# A comprehensive survey of genetic variation in 20,691 subjects from four large cohorts

**DOI:** 10.1101/083030

**Authors:** Sara Lindström, Stephanie Loomis, Constance Turman, Hongyan Huang, Jinyan Huang, Hugues Aschard, Andrew T. Chan, Hyon Choi, Marilyn Cornelis, Gary Curhan, Immaculata De Vivo, A. Heather Eliassen, Charles Fuchs, Michael Gaziano, Susan E. Hankinson, Frank Hu, Majken Jensen, Jae H. Kang, Christopher Kabrhel, Liming Liang, Louis R. Pasquale, Eric Rimm, Meir J. Stampfer, Rulla M. Tamimi, Shelley S. Tworoger, Janey L. Wiggs, David J. Hunter, Peter Kraft

## Abstract

The Nurses’ Health Study (NHS), Nurses’ Health Study II (NHSII), Health Professionals Follow Up Study (HPFS) and the Physicians Health Study (PHS) have collected detailed longitudinal data on multiple exposures and traits for approximately 310,000 study participants over the last 35 years. Over 160,000 study participants across the cohorts have donated a DNA sample and to date, 20,691 subjects have been genotyped as part of genome-wide association studies (GWAS) of twelve primary outcomes. However, these studies utilized six different GWAS arrays making it difficult to conduct analyses of secondary phenotypes or share controls across studies. To allow for secondary analyses of these data, we have created three new datasets merged by platform family and performed imputation using a common reference panel, the 1,000 Genomes Phase I release. Here, we describe the methodology behind the data merging and imputation and present imputation quality statistics and association results from two GWAS of secondary phenotypes (body mass index (BMI) and venous thromboembolism (VTE)).

We observed the strongest BMI association for the FTO SNP rs55872725 (β=0.45, p=3.48×10^−22^), and using a significance level of p=0.05, we replicated 19 out of 32 known BMI SNPs. For VTE, we observed the strongest association for the rs2040445 SNP (OR=2.17, 95% CI: 1.79-2.63, p=2.70×10^−15^), located downstream of F5 and also observed significant associations for the known ABO and F11 regions. This pooled resource can be used to maximize power in GWAS of phenotypes collected across the cohorts and for studying gene-environment interactions as well as rare phenotypes and genotypes.

## INTRODUCTION

Large, well-phenotyped cohort studies have constituted the backbone of epidemiology for several decades. Prospectively collected longitudinal information on exposures and outcomes enables a broad spectrum of analyses and has led to novel insights into disease etiology, such as the link between smoking and lung cancer [1,2] as well as the link between both high cholesterol levels and trans fatty acids with coronary heart disease [3,4] Many existing cohorts collect biological specimens from their participants, allowing for studies of inherited genetic variation as well as prospectively measured biomarkers such as metabolomic profiles [5] and circulating hormone levels [6]. Genome-wide association studies (GWAS) are currently a main engine of genetic epidemiology and have led to the identification of thousands of loci for hundreds of traits (for an overview and its clinical applications, see Manolio [7]). When designing a GWAS, cost is still the determining factor and consequently, GWAS within cohorts are often conducted within nested case-control studies or sub-cohorts. In contrast, the Women’s Genome Health Study (WGHS) [8] genotyped the entire cohort of 27,000 women and the Genetic Epidemiology Research on Adult Health and Aging (GERA) Cohort has generated GWAS data on almost 100,000 individuals [9]. However, in many instances, GWAS are tied to specific funding sources acquired for studying a pre-defined outcome and only a small fraction of the cohort is genotyped at a specific time.

Within the Nurses’ Health Study (NHS) [10], Nurses’ Health Study II (NHSII) [11], Health Professional Follow Up Study (HPFS) [12] and the Physicians’ Health Study (PHS) [13], since 2007, we have, conducted twelve GWAS of different traits including type 2 diabetes [14], coronary heart disease [15], several cancer types [16-19] and mammographic density [20,21]. In total, we have assembled GWAS data for 20,769 individuals across the cohorts, creating unprecedented opportunities to conduct secondary analyses on other collected outcomes. Indeed, we have used one or many of these GWAS to analyze secondary phenotypes including but not limited to body anthropometrics [22-24], hair color [25], reproductive aging [26], smoking behavior [27], telomere length [28], mammographic density [29], cutaneous nevi [30], melanoma [30], depressive symptoms [31], coffee consumption [32] as well as circulating levels of B12 [33], folate [34], hormones [35], vitamins [36,37], retinol [38] and e-selectin [39]. However, GWAS of secondary traits face practical issues in terms of different genotyping arrays, low variability in the phenotype of interest within a single GWAS (e.g. rare diseases where only a handful of cases may occur in the original GWAS), and theoretical issues including ascertainment bias due to oversampling of cases [40] or differential genotype/imputation quality between studies [41] (e.g. if controls are “utilized” from GWAS data generated on a different genotype platform).

Here, we describe our pipeline for merging and imputing the individual GWAS datasets within NHS, NHSII, HPFS and PHS. Datasets were merged based on genotype platform family and all data were subsequently imputed to a common reference panel (the 1,000 Genomes Phase I release [42]). We present proof-of-principle results from genome-wide analysis of body mass index (BMI) and venous thromboembolism (VTE).

## METHODS

### Description of NHS, NHSII, HPFS and PHS

In 1976, the Nurses’ Health Study (NHS) was launched with the goal of studying women’s health [10]. Since that time, 121,700 nurse participants have answered biennial questionnaires (response rate >90% over time) about personal and physical characteristics, physical activity and ability, reproductive history, family history of disease, environmental/personal exposures, diet and dietary supplements, screening, disease and health conditions, prescription and over-the-counter medications, and psychosocial history. In addition, 32,826 blood and 29,684 cheek cell samples have been collected since the late 1980s. An additional 116,430 nurses were recruited in 1989 as a part of Nurses’ Health Study II (NHSII) and have returned biennial questionnaires similar to those used for NHS [11]. For NHSII, we have collected blood samples for 29,612 women and cheek cell samples for an additional 29,859 women. The Health Professional Follow-Up Study (HPFS) began in 1986 with the aim of studying men's health [12]. A total of 51,529 men in health professions were recruited, and every two years, members of the study receive questionnaires similar to the ones used in NHS. In HPFS, we have collected blood samples from 18,159 participants and cheek cell samples from an additional 13,956 men. The Physicians’ Health Study (PHS) is a randomized primary prevention trial of aspirin and supplements among 29,067 United States physicians followed with annual questionnaires since 1982 [13]. A total of 14,916 men provided a baseline blood sample.

### Ethics Statement

Each GWAS study was approved by the Brigham and Women’s Hospital Institutional Review Board. Return of the mailed self-administered questionnaires was voluntary. Thus, receipt of a completed questionnaire was considered as evidence of a desire to participate in the study and was taken as a formal indication of consent.

### Description of GWAS studies and genotyping

Since 2007, twelve separate GWAS have been conducted within these four cohorts **(Table 1).** The primary traits are breast cancer [16], pancreatic cancer [43], glaucoma [44], endometrial cancer [17], colon cancer [19], glioma [45], prostate cancer [18], type 2 diabetes [14], coronary heart disease [15], kidney stones, gout and mammographic density [20]. These studies were genotyped on six different arrays **(Table 1)** at four different genotyping centers (National Cancer Institute, Broad Institute, University of Southern California and Rosetta/Merck). Standard quality control filters for call rate, Hardy-Weinberg equilibrium, and other measures were applied to the genotyped SNPs and/or samples. In total, these GWAS data sets comprise 20,769 participants including 11,522 from NHS, 934 subjects from NHSII, 7,018 subjects from HPFS and 1,305 subjects from PHS.

**Table 1:** **GWAS datasets in HPFS, NHS, NHSII and PHS**

| Cohort | Outcome | Subjects<br>(cases/controls) | Platform | GWAS dataset |
| --- | --- | --- | --- | --- |
| HPFS | Coronary Heart Disease | 435/878 | Affymetrix 6.0 | AffyMetrix |
| HPFS | Type 2 Diabetes | 1,189/1,298 | Affymetrix 6.0 | AffyMetrix |
| HPFS | Pancreatic Cancer | 54/52 | Illumina 550k | Illumina HumanHap |
| HPFS | Kidney Stone | 315/238 | Illumina 610k | Illumina HumanHap |
| HPFS | Prostate Cancer | 218/205 | Illumina 610k | Illumina HumanHap |
| HPFS | Glaucoma | 178/299 | Illumina 660W | Illumina HumanHap |
| HPFS | Glioma | 26/0 | Illumina 660W | Illumina HumanHap |
| HPFS | Colon Cancer | 229/230 | Illumina OmniExpress | Illumina OmniExpress |
| HPFS | Gout | 717/699 | Illumina OmniExpress | Illumina OmniExpress |
|  | **SUBTOTAL** |  |  | **7,018 (1,511 Illumina Human Hapmap, 3,634 Affymetrix, 1,873 Illumina OmniExpress)** |
| NHS | Type 2 Diabetes | 1,532/1,754 | Affymetrix 6.0 | AffyMetrix |
| NHS | Coronary Heart Disease | 342/804 | Affymetrix 6.0 | AffyMetrix |
| NHS | Ovarian Cancer | 36/0 | Illumina 317k | Illumina HumanHap |
| NHS | Breast Cancer | 1,145/1,142 | Illumina 550k | Illumina HumanHap |
| NHS | Pancreatic Cancer | 82/84 | Illumina 550k | Illumina HumanHap |
| NHS | Kidney Stone | 328/166 | Illumina 610k | Illumina HumanHap |
| NHS | Glaucoma | 313/497 | Illumina 660W | Illumina HumanHap |
| NHS | Glioma | 38/0 | Illumina 660W | Illumina HumanHap |
| NHS | Endometrial Cancer | 396/348 | Illumina OmniExpress | Illumina OmniExpress |
| NHS | Colon Cancer | 394/774 | Illumina OmniExpress | Illumina OmniExpress |
| NHS | Mammographic density | 153/641 | Illumina OmniExpress | Illumina OmniExpress |
| NHS | Gout | 319/392 | Illumina OmniExpress | Illumina OmniExpress |
|  | **SUBTOTAL** |  |  | **11,522 (3,711 Illumina Human Hapmap, 4,413 Affymetrix, 3,380 Illumina OmniExpress)** |
| NHSII | Breast Cancer | 289/0 | Illumina 610k | Illumina HumanHap |
| NHSII | Kidney Stone | 341/294 | Illumina 610k | Illumina HumanHap |
|  | **SUBTOTAL** |  |  | **924 (924 Illumina Human Hapmap, 0 Affymetrix, 0 Illumina OmniExpress)** |
| PHS | Pancreatic Cancer | 49/54 | Illumina 550k | Illumina HumanHap |
| PHS | Prostate Cancer | 312/363 | Illumina 610k | Illumina HumanHap |
| PHS | Colon Cancer | 331/333 | Illumina OmniExpress | Illumina OmniExpress |
|  | **SUBTOTAL** |  |  | **1,305 (641 Illumina Human Hapmap, 0 Affymetrix, 664 Illumina OmniExpress)** |
|  | **TOTAL** |  |  | **20,769 (6,787 Illumina Human Hapmap, 8,065 Affymetrix, 5,917 Illumina OmniExpress)** |

### Dataset merging

Successfully merging genotype data for different individuals requires complete overlap in SNPs. SNPs that are missing by design (due to different genotyping platforms) from some studies will be correlated with the primary phenotype for that dataset. This might cause spurious results in any secondary analysis on related traits. Although a missing SNP can be imputed, it will have a higher degree of inaccuracy in imputed compared with genotyped SNPs, potentially creating differential measurement error that could also lead to bias [41,46,47]. Therefore, we first looked at the overlap of SNPs between different genotyping arrays and identified three broad platform families with high degree of overlap within category but low overlap across categories – the earlier generation of Illumina arrays (HumanHap), the Illumina OmniExpress array and Affymetrix 6.0 array. The HumanHap platform had a total of 459,999 SNPs compared with 565,810 SNPs for OmniExpress and 668,283 SNPs for Affymetrix 6.0. However, the intersection among all three platform families was only 75,285 SNPs **(Figure 1).** To achieve the largest GWAS datasets as possible without losing SNP information, we created three datasets – HumanHap comprising six GWAS datasets, OmniExpress comprising four GWAS datasets and Affymetrix 6.0 comprising two GWAS datasets. In the merging process, we removed any SNPs that were not in all studies for a specific platform or had a missing call rate>5%. We flipped strands where appropriate and removed A/T and C/G SNPs to create the final compiled datasets.

**Figure 1:** Overlap in SNPs across genotype platforms

We ran a pairwise identity by descent (IBD) analysis within and across the combined dataset to detect duplicate and related individuals based on resulting IBD probabilities Z_0_, Z_1_ and Z_2_ (Z_k_ is probability that a pair of subjects share k alleles identical by descent, estimated from genome-wide SNP data). If 0≤Z_0_≤0.1 and 0≤Z_1_≤0.1 and 0.9≤Z_2_≤1.1 then a pair was flagged as being identical twins or duplicates. Pairs were considered full siblings if 0.17≤Z_0_≤0.33 and 0.4≤Z_1_≤0.6 and 0.17≤Z_2_≤0.33. Half siblings or avunculars were defined as having 0.4≤Z_1_≤0.6 and 0≤Z_2_≤0.1. Some of the duplicates flagged were expected, having been genotyped in multiple datasets and hence having the same cohort identifiers. In this case, one of each pair was randomly chosen for removal from the dataset. In instances where pairs showed pairwise genotype concordance rate>0.999 but were not expected duplicates, both individuals were removed. Related individuals (full siblings, half siblings/avunculars) were not removed from the final datasets. In the HumanHap dataset, 107 individuals were removed because they were duplicates or flagged for removal in the genotyping step, leaving 6,787 subjects. In addition, 8 pairs of individuals were flagged as related. In the OmniExpress dataset, we removed 39 subjects leaving 5,917 IDs and 5 pairs of related subjects. In the Affymetrix dataset, 167 individuals were removed because they were duplicates or were flagged for removal from secondary genotype data cleaning, leaving a total of 8,065 individuals. Across all three datasets, we identified 444 duplicate pairs (406 expected) and thus removed additional 482 individuals from analysis across all three platform families.

After removing duplicate and related pairs of IDs, we used EIGENSTRAT [48] to run principal component analysis (PCA) on each dataset, removing one member from each flagged pair of related individuals. For Affymetrix and HumanHap, we used approximately 12,000 SNPs from Yu et al [49] that were filtered to ensure low pairwise linkage disequilibrium (LD). For the OmniExpress dataset we used approximately 33,000 SNPs that were similarly filtered. The top principal components were manually checked for outliers.

To identify any SNPs that created spurious associations, we ran several logistic regression analyses among subjects that were selected as controls in the initial GWAS (i.e. excluding all case subjects). For each regression, we used cohort-specific controls from one original GWAS as cases and the rest of the controls in that dataset as controls. For example, in the OmniExpress dataset, we considered NHS controls from the gout GWAS as “cases” while treating controls from the gout (HPFS), endometrial cancer (NHS), colon cancer (NHS, HPFS and PHS), and mammographic density (NHS) as “controls”. We repeated this, treating each cohort-specific “controls set” as “cases” and all other controls as “controls”. For each GWAS, we extracted genome-wide significant SNPs (p<10^−8^) and examined QQ plots. In the Affymetrix dataset, 100 SNPs were flagged and removed. In the HumanHap dataset, 8 SNPs had p<10^−8^ in at least one of the QC regressions and were removed. No SNPs in the OmniExpress dataset had p<10^−8^ and hence, no SNP was removed.

### Imputation

After the datasets were combined and appropriate SNP and subjects filters applied, the compiled datasets were separately imputed. We used the 1000 Genomes Project ALL Phase I Integrated Release Version 3 Haplotypes excluding monomorphic and singleton sites (2010-11 data freeze, 2012-03-14 haplotypes) as the reference panel. SNP and indel genotypes were imputed in three steps. First, genotypes on each chromosome were split into chunks to facilitate windowed imputation in parallel using ChunkChromosome (v.2011-08-05). Then each chunk of chromosome was phased using MACH [50,51] (v.1.0.18.c). In the final step, Minimac (v.2012-08-15) was used to impute the phased genotypes to approximately 31 million markers in the 1000 Genomes Project.

### “Proof of Principle” GWAS– BMI and VTE

To validate our merged GWAS datasets, we conducted two proof-of principle GWAS of one quantitative trait (BMI) and one binary trait (VTE). We defined BMI as weight (kg)/height^2^ (cm) and obtained it by extracting information on weight from the accompanying questionnaire collected at time of blood draw. If weight information was missing, we extracted it from the questionnaire closest in time to time of blood draw. Height was extracted from the baseline questionnaire. We obtained data on BMI for 20,283 participants. VTE is a spectrum of disease that includes pulmonary embolism (PE) and deep vein thromboembolism (DVT). Physician-diagnosed PE has been asked on every biennial NHS questionnaire since 1982, and every NHSII and HPFS questionnaire since cohort inception. In the NHS, DVT without PE is captured when a nurse answers that she has had phlebitis or thrombophlebitis (ICD-9=453.x). In NHS, NHSII and HPFS cohorts through 2010 (we did not have VTE data for PHS), we identified 6,041 individuals who reported VTE. Self-reported PE was verified through medical records review by a trained physician (CK). DVT cases are based on self-report, though a validation study of 100 DVT cases found self-reports to be highly consistent (>96%) with medical record review. In total, we identified 1,364 VTE cases with GWAS data. We treated all non-VTE cases with GWAS data as controls (n=17,628). Since we did not have data on VTE in PHS, we excluded PHS from this analysis.

### Statistical analysis – GWAS

SNPs and indels with an imputation quality score <0.3 (as defined by the RSQR_HAT value in MACH) or a minor allele frequency (MAF) <0.01 were excluded. Primary association analysis was performed separately within each platform family (HumanHap, OmniExpress and Affymetrix). For imputed SNPs, the estimated number of effect alleles (ranging from 0 to 2) was used as a covariate. For BMI, we conducted linear regression adjusting for study (indicator variables including cohort as well as primary GWAS outcome), age at blood draw and the top four principal components. For VTE, we conducted logistic regression adjusting for study as above and the top four principal components. For both BMI and VTE, we combined platform family-specific results with fixed-effects meta-analysis using the METAL [52] software. We used the Cochran’s Q statistic to test for heterogeneity across studies.

## RESULTS

### Imputation statistics

We imputed a total of 31,326,389 markers (29,890,747 SNPs and 1,435,642 indels) and the majority (69%) of these had a MAF<0.01. The average imputation quality score by minor frequency for each platform family is shown in **Figure 2.** The imputation quality was very similar across all three datasets **(Suppl Figures 1-3)** with 49-51% of markers having an imputation quality score ≥0.3. When restricting to markers with MAF>0.01 (~10 million), 92-94% of the markers had a quality score ≥0.3. After filtering markers based on MAF (>0.01) and imputation r-sq (≥0.3), approximately 9.8 million markers were available for analysis.

**Figure 2:** Imputation quality score by minor allele frequency for the three platform families

### BMI results

We had BMI and GWAS data for 20,283 individuals (n=6,762 for HumanHap, n=5,844 for OmniExpress, n=7,677 for AffyMetrix) within NHS, NHSII, HPFS and PHS. Platform-specific QQ-plots **(Suppl Figures 4a-c)** showed no indication of systematic bias (genomic inflation factor λ=1.00-1.02). The results from the meta-analysis are shown in **Figures 3 and 4.** We observed a tail of strongly associated SNPs with the top SNPs located in the known BMI *FTO* locus (strongest associated SNP: rs55872725, β=0.45, p=3.48×10^−22^). We also observed genome-wide significant associations for the previously identified *TMEM18* (strongest associated SNP: rs7563362, β=−0.36, p=1.76×10^−8^) and *FANCL* loci (strongest associated SNP: rs980183, β=−0.26, p=2.73×10^−8^). Using a significance level of p=0.05, 59% (19/32) known BMI SNPs [53], showed association with BMI in our data. In addition, 31 out of the 32 known SNPs showed associations in the same direction as the original BMI study **(Figure 5).**

**Figure 3:** QQ-plot for GWAS analysis of body mass index based on 20,283 individuals.

**Figure 4:** Manhattan plot for GWAS analysis of body mass index based on 20,283 individuals.

**Figure 5:** Associations for known body mass index.

### VTE results

We had information on VTE status and GWAS data for 1,364 cases and 17,628 controls within NHS, NHSII and HPFS. The median number of case subjects by dataset was 87.5 and ranged from 16 in the NHSII breast cancer GWAS dataset (total of 289 individuals) to 417 in the type 2 diabetes GWAS dataset (total of 5,773 individuals). The small number of cases in many individual GWAS data sets led to unstable study-specific association statistics. Restricting to studies with an expected case minor allele count >10 for SNPs with a MAF of 0.05 (i.e. studies with at least 200 cases) reduced the sample size to 417 cases and 5,356 controls. However, within each compiled imputed GWAS dataset, VTE case numbers ranged from 406 (OmniExpress) to 532 (Affymetrix). Thus, combining the individual GWAS datasets into three main datasets enabled association analysis of hundreds of cases rather than tens, leading to more stable estimates in the regression analysis. Platform-specific QQ-plots **(Suppl Figures 5a-c)** showed no indication of systematic bias (genomic inflation factor λ=1.00-1.01). The results from the meta-analysis are shown in **Figures 6 and 7** (genomic inflation factor λ=1.00). We observed a strong association located downstream of the *F5* gene (strongest associated SNP: rs2040445, OR=2.17, 95% CI: 1.79-2.63, p=2.70×10^−15^). We also observed genome-wide significant associations for the ABO locus (strongest associated SNP: rs2519093, OR=1.36, 95% CI: 1.23-1.49, p=1.51×10^−10^) and a nominal association (P=0.007) with the previously VTE-associated *F11* locus.

**Figure 6:** QQ-plot for GWAS analysis of venous thromboembolism based on 1,364 cases and 17,628 controls.

**Figure 7:** Manhattan plot for GWAS analysis of venous thromboembolism based on 1,364 cases and 17,628 controls.

## DISCUSSION

Thousands of genetic loci associated with hundreds of complex traits have been identified through GWAS and as sample sizes continue to increase, more loci will be discovered. Although the cost of GWAS has dropped, lack of financial resources is still the limiting factor for generating new data. Most GWAS have been conducted in case-control studies, and this has led to the creation of disease-specific consortia in which power can be maximized. However, there is usually only one disease phenotype available from these cases, and little capacity to follow cases or controls to collect information on additional phenotypes that develop over time. Cohort studies are designed to collect multiple endpoints on individuals, but often suffer from limited power for a specific disease. To maximize the utility of existing cohort data resources, it is important to explore associations with additional traits and outcomes that have been collected for individuals in multiple cohorts. In particular, the accumulation of GWAS data within large cohorts with rich environmental and outcome data creates new opportunities to assess novel hypotheses. In addition, cohort studies provide unique opportunities to prospectively assess biomarker-disease associations, thereby minimizing bias due to reverse causation or treatment effects. However, “borrowing” GWAS data between traits is not straightforward. Known issues that can cause bias include technical artifacts due to different genotyping platforms, differences in imputation accuracy and ascertainment bias. Thus, careful data management, imputation procedures and quality checks are needed. Furthermore, if the secondary trait is rare, there will be low phenotypic variability within each GWAS dataset. For example, we observed fewer than 100 VTE cases within the majority of individual GWAS, compared to more than 400 cases within each combined dataset.

Our pipeline for combining and imputing twelve different GWAS datasets can overcome both technical and methodological issues. We chose to create three different datasets defined by platform family (in our case, Illumina HumanHap, Illumina OmniExpress and AffyMetrix) since the SNP overlap across platforms was low on a genome-wide scale (75,285 SNPs). An attempt to impute a genome-wide dataset comprising only 75,000 SNPs as starting point would have resulted in decreased imputation accuracy in regions of the genome with sparse genotype data. Moreover, it has been shown that different platforms might call SNPs differently and that SNP-specific allele frequencies can differ between platforms (see [41] for further discussion). We conducted multiple case-control GWAS among control subjects within each dataset (i.e. running multiple “null” GWAS) and identified and excluded more than 100 SNPs that showed spurious associations. These results emphasize that although datasets are merged by platform family, problematic SNPs giving rise to spurious associations might still exist and it is important to carefully check for these.

To assess the validity of our data, we conducted two proof-of-principle GWAS. The first trait we studied was BMI, and in line with what expected, we observed strong evidence of associations with known BMI loci including *FTO* and *TMEM18* that both reached genome-wide significance (P<5×10^−8^). In addition, out of 32 known BMI SNPs we observed nominal significance (P<0.05) for 19 of them, all in the same direction as expected from previous reports. Of note, our sample size (n=20,823) is less than 10% of the original GWAS that had a total sample size of 249,766 individuals. Therefore, we would not expect to observe significant associations for all BMI SNPs due to limited power. For VTE, we observed genome-wide significant associations for the *F5* and *ABO* loci that are both known to be associated with VTE. In addition, we also observed a nominal association (P=0.007) with the *F11* region. Our BMI and VTE results confirm that GWAS analysis of secondary traits in this data is valid and provides a platform for future studies of secondary traits. We ran the BMI and VTE analyses twice, the first time without removing duplicates between the datasets (total of 444 pairs), and the second time with the duplicates removed. Although the 444 pairs constitute less than 5% of our total sample size, including them had an impact on the genomic inflation factor (for BMI, the genomic inflation factor went from 1.09 to 1.05 and for VTE, the genomic inflation factor went from 1.02 to 1.00). These results are especially interesting as it is often difficult to identify duplicates across studies when raw data from all participating studies are not available. Care should be taken to remove overlapping subjects across GWAS contributing to a meta-analysis, but any remaining cryptic overlap may inflate association statistics. In that case, statistical adjustment procedures like LD score regression [54] can be used to account for cryptic overlap.

One of the main benefits with collecting comprehensive genetic information on cohort subjects is the opportunity to assess interactions between genetic factors and prospectively collected environmental data. To date, few gene-environment interactions have been identified and although their extent and clinical impact remain an open empirical question, the current lack of homogenous large datasets with both genetic and environmental data has precluded comprehensive investigation. Capitalizing on this GWAS resource, we will be able to explore gene-environment interactions for a plethora of outcomes including complex traits such as height and BMI, but also disease outcomes. It will also allow us to study the impact of environmental factors within genetic strata to identify individuals for whom a particular intervention might be especially important [55-58].

Accumulation of these GWAS data is ongoing and we expect to generate new GWAS data for an additional 15,000 participants within the next two years, almost doubling our total GWAS sample size. This growing resource will be a core component of future studies aiming to elucidate how genes and the environment impact public health.

## ACKNOWLEDGEMENTS

This work was supported by National Institute of Health (P01CA87969, P01CA055075, P01DK070756, U01HG004728, UM1CA186107, UM1CA176726, R01CA49449, R01CA50385, R01CA67262, R01CA131332, R01HL034594, R01HL088521, R01HL35464, R01HL116854, R01EY015473, R01EY022305, P30EY014104, R03DC013373 and R03CA165131). Dr. Pasquale is also supported by a Harvard Medical School Distinguished Ophthalmology Scholar award. The funders had no role in study design, data collection and analysis, decision to publish, or preparation of the manuscript.

---

**Supplemental Figure 1:** Proportion of sucessfully imputed markers on the Affymetrix platform

**Supplemental Figure 2:** Proportion of sucessfully imputed markers on the Illumina HumanHap platform.

**Supplemental Figure 3:** Proportion of sucessfully imputed markers on the Illumina Omniexpress platform.

**Supplemental Figure 4a:** QQ-plot for GWAS analysis of body mass index on the Illumina Omniexpress platform (n=5,844).

**Supplemental Figure 4b:** QQ-plot for GWAS analysis of body mass index on the Affymetrix platform (n=7,677).

**Supplemental Figure 4c:** QQ-plot for GWAS analysis of body mass index on the Illumina HumanHap platform (n=6,762).

**Supplemental Figure 5a:** QQ-plot for GWAS analysis of venous on the Illumina Omniexpress platform (406 cases and 4,786 controls).

**Supplemental Figure 5b:** QQ-plot for GWAS analysis of venous on the Illumina Omniexpress platform (406 cases and 4,786 controls).

**Supplemental Figure 5c:** QQ-plot for GWAS analysis of venous on the Affymetrix platform (532 cases and 7,147 controls).

